# Rethomics: an R framework to analyse high-throughput behavioural data

**DOI:** 10.1101/305664

**Authors:** Quentin Geissmann, Luis Garcia Rodriguez, Esteban J. Beckwith, Giorgio F. Gilestro

## Abstract

The recent development of automatised methods to score various behaviours on a large number of animals provides biologists with an unprecedented set of tools to decipher these complex phenotypes. Analysing such data comes with several challenges that are largely shared across acquisition platform and paradigms. Here, we present `rethomics`, a set of `R` packages that unifies the analysis of behavioural datasets in an efficient and flexible manner. `rethomics` offers a computational solution to storing, manipulating and visualising large amounts of behavioural data. We propose it as a tool to bridge the gap between behavioural biology and data sciences, thus connecting computational and behavioural scientists. `rethomics` comes with a extensive documentation as well as a set of both practical and theoretical tutorials (available at https://rethomics.github.io).

## Introduction

The biological implications and the determinism of animal behaviour have long been a subject of significant scientific interest. In the 1960s, behavioural geneticists, such as Seymour Benzer, showed that some apparently complex behaviours could in fact be governed by simple genetic determinants, which connected genetics and behaviour [1]. In the last few decades, our ability to acquire and analyse vast amounts of biological data has tremendously increased [2], deeply transforming both genetics [3] and neuroscience [4]. In fact, ethology itself is undergoing its own transition towards data sciences, which has prompted the terms ‘ethomics’ [5,6] and ‘computational ethology’ [7]. It is now accepted that the study of behaviour can also benefit from quantitative sciences such as machine learning, physics and computational linguistics [8,9].

Although general questions regarding the environmental, evolutionary, neural and genetic determinants of behaviours are shared within the community, the multiplicity of model organisms, hypotheses and paradigms has led to the existence of a very diverse palette of specific recording techniques. Some tools were developed, for instance, to continuously record simple behavioural features such as walking activity [10] and position [11] over long durations (days or weeks); to score more complex ones such as feeding [12,13], aggression [14] and courtship [15]; and to study the behaviour of multiple interacting animals [16–18]. Whilst most recording platforms are unrelated to each other, there are also some attempts to build general purpose tools that can be adapted by researchers to suit their specific goals [5,19–21]. However, when it comes to the subsequent analysis of the generated results, there is still no unified programmatic framework that could be used as a set of building blocks in a pipeline.

The fields of structural biology and bioinformatics are good examples of communities that have taken advantage of sharing standard files formats, modular command line tools [22] and software packages [23] that can be assembled into pipelines [24]. In these research areas, which are closely linked to data sciences and statistics, scripting interfaces are the standard since they help to deliver reproducible results [25,26]. In addition, they can be used on remote resources such as computer clusters, which makes them more scalable to the context of ‘big data’ [27]. Since many aspects of behaviour analysis are also becoming increasingly dependent on data sciences, the development of such common tools and data structures would be very valuable.

At first, it may seem as though behavioural experiments are prohibitively heterogeneous – in terms of model organisms, paradigm and timescale – for a similar community to arise. However, low-level conceptual consistencies and methodological challenges are shared across experiments. For instance, the results (*i.e.* the ‘data’) feature a set of long time series (sometimes multivariate and irregular), but also contain a formal description of the treatment applied to each individual, the ‘metadata’. Storing and accessing data and metadata efficiently involves the implementation of a nested data structure which, in principle, can be shared between acquisition platforms and experimental paradigms.

Here, we describe the `rethomics` platform, an effort to promote the interaction between behavioural biologists and data scientists. `rethomics` is implemented as a collection of interconnected packages, offering solutions to importing, storing, manipulating and visualising large amounts of behavioural results. We also present two practical examples of its application to the analysis of behavioural rhythm in fruit flies, a widely studied subject.

## Design and Implementation

`rethomics` is implemented in `R` [28] as a collection of packages (Fig 1). Such modular architecture follows the model of modern frameworks such as the `tidyverse` [29], which results in increased testability, maintainability and adaptability. In this model, each task of the analysis workflow (*i.e.* data import, manipulation and visualisation) is handled by a different package, and new ones can be designed to suit specific needs. At the core of `rethomics` lies the `behavr` table, a structure used to store large amounts of data (*e.g.* position and activity) and metadata (*e.g.* treatment and genotype) in a unique `data.table`-derived object [30]. Any input package will import experimental data as a `behavr` table which can, in turn, be analysed and visualised regardless of the original input platform. Numerical results and plots are standard objects that can, therefore, be further analysed inside the wide `R` package ecosystem.

**Fig 1.**
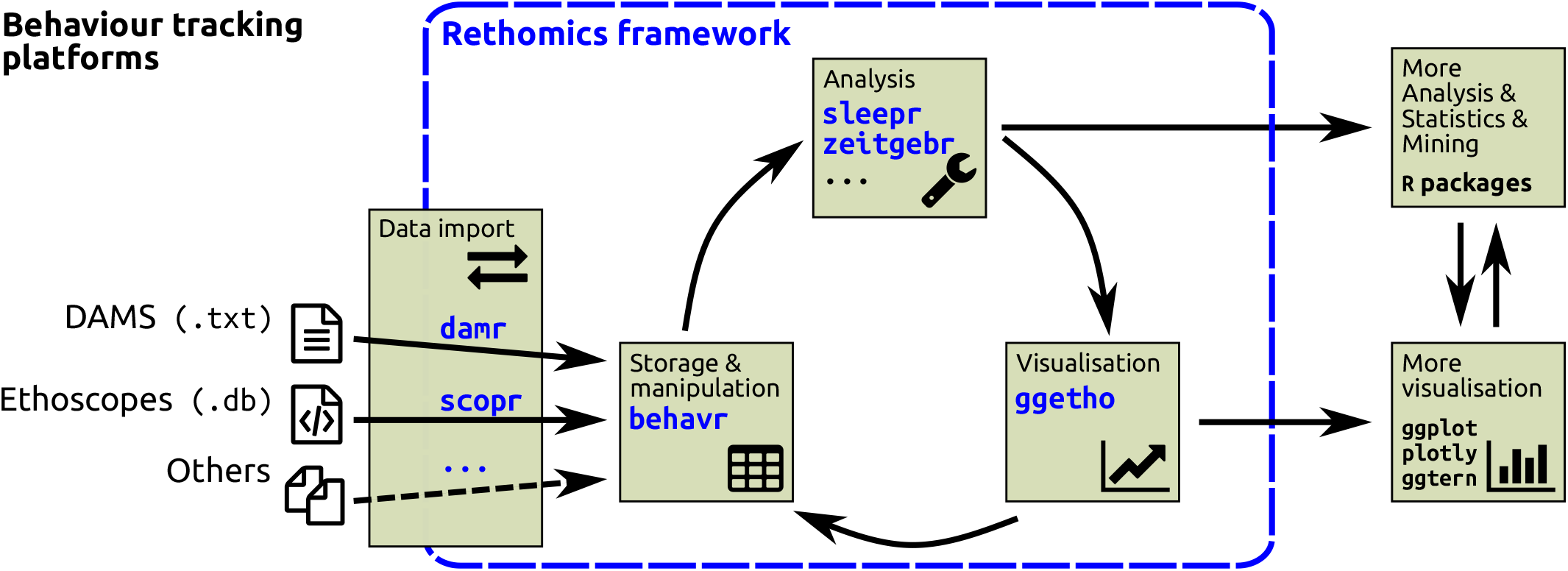
The rethomics workflow. Diagram representing the interplay between, from left to right, the raw data, the `rethomics` packages (in blue) and the rest of the `R` ecosystem.

### Internal data structure

Ethomics results can easily scale and data structure therefore gains from being computationally efficient – both in terms of memory footprint and processing speed. For instance, there could be very long time series, sampled several times per second, over multiple days, for each individual. In addition, time series can be multivariate – encoding coordinates, orientation, dimensions, activity, colour intensity and so on. Furthermore, experiments may feature a large number of individuals. Each individual is also associated with some metadata: a set of ‘metavariables’ that describe experimental conditions. For instance, metadata stores information regarding the date and location of the experiment, treatment, genotype, sex, *post hoc* observations and other arbitrary metavariables. A large set of metavariables is an important asset since they can later be treated as covariates.

`behavr` tables link metadata and data within the same object, extending the syntax of `data.table` to manipulate, join and access metadata (Fig 2 and S1 Fig). This approach guarantees that any data point can be mapped correctly to its parent metadata thanks to a shared key (`id`). Furthermore, it allows implicit update of metadata when data is altered. For instance, when data is subsetted, only the remaining individuals should be in the new metadata. It is also important that metadata and data can interoperate – for example, when updating a variable according to the value of a metavariable (say, altering the variable x only for animals with the metavariable `sex = ‘male’`). The online tutorials and documentation provide a detailed set of examples and concrete use cases of `behavr`.

**Fig 2.**
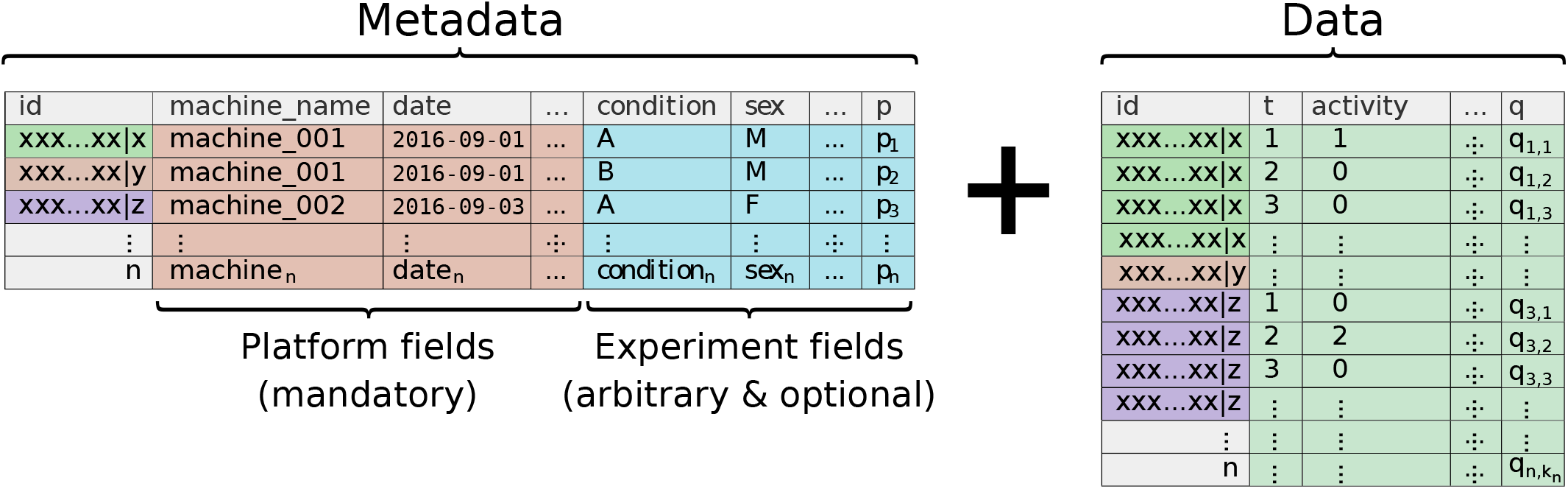
behavr table. Illustration of a `behavr` object, the core data structure in `rethomics`. The metadata holds a single row for each of the *n* individuals. Its columns, the *p* metavariables, are one of two kinds: either required – and defined by the acquisition platform (*i.e.* used to fetch the data) – or user-defined (*i.e.* arbitrary). In the data, each row is a ‘read’ (*i.e.* information about one individual at one time-point). It is formed of *q* variables and is expected to have a very large number of reads, *k*, for each individual *i* ∈ [1, *n*]. Data and metadata are implicitly joined on the `id` field. Note that the names used for variables and metavariable in this example are only plausible cases which will likely differ in practice.

### Data import

Data import packages translate results from a specific recording platform (*e.g.* text files and relational databases) into a single `behavr` object. Currently, we provide two packages: one to import data from single and multi-beam Drosophila Activity Monitor Systems (Trikinetics Inc.) and another one for Ethoscopes [21]. Although the structure of the raw results is very different, conceptually, loading data is very similar. In all cases, users must provide a metadata table, with one row per individual, and featuring both mandatory and optional columns (Fig 2). The mandatory ones are the necessary and sufficient information to fetch data (*e.g.* machine id, region of interest and date). The optional columns are user-defined arbitrary fields that translate experimental conditions (*e.g.* treatment, genotype and sex).

In this respect, the metadata file is a standardised and comprehensive data frame describing an experiment. It explicitly lists all treatments and individuals, which facilitates interspersion of conditions. Furthermore, it streamlines the inclusion and analysis of further replicates in the same workflow. Indeed, additional replicates can simply be added as new rows – and the `id` of the replicate later used, if needed, as a covariate.

### Visualisation

To integrate visualisation in `rethomics`, we implemented `ggetho`, a package that offers new tools that extend the widely adopted `ggplot2` [31]. `ggetho` makes full use of the internal `behavr` structure to summarise temporal trends. We implemented a set of new ‘layers’ and ‘scales’ that particularly applies to the visualisation of long experiments, with the ability to, for instance, display ‘double-plotted actograms’, periodograms, annotate light and dark phases and wrap time over a given period. Importantly, `ggetho` is fully compatible with `ggplot2`. For instance, `ggplot2` operations such as faceting, transforming axes and adding new layers functions natively with `ggetho`.

## Results

In order to illustrate the usefulness of `rethomics`, we provide two examples. The first one is a detailed and reproducible description of the loading and analysis activity data in the context of circadian rhythm, using DAM2 (Trikinetics Inc.) data. The second one shows how `rethomics` integrates within `R` to perform a multi-scale analysis of a periodic behaviour, using continuous wavelet transform, on data generated with ethoscopes [21].

### Canonical circadian analysis in *Drosophila*

The `zeitgebr` package implements a comprehensive suite of methods to analyse circadian rhythms, including the computation of autocorrelograms, *χ*^2^ [32] and Lomb-Scargle [33] periodograms, and automatic peak detection.

The study of the rhythmical activity of fruit flies has played a crucial role in the development of circadian biology [34]. To date, most of the behavioural data used in the field is acquired through the Drosophila Activity Monitor System (DAMS). The package `damr` was developed to import DAMS results in the `rethomics` framework, which we envision will be a common use case. To illustrate the ability of `rethomics` to analyse pre-existing results, we gathered a subset of the data from a recent publication [35], kindly made publicly available by the authors [36].

Wild type flies are highly rhythmic in Light-Dark (LD) cycles and become arrhythmic in constant light (LL). In their study, the authors gain understanding of the function of the molecular clock by showing that overexpression of the gene NKCC makes the flies rhythmic in LL, and that the endogenous period in LL is longer than 24 hours.

Here, we guide the reader through the analysis of two of the genotypes employed in that study; one control group (NKCC^ox^/+, which is arrhythmic under LL) and one where NKCC^ox^ is overexpressed in clock neurons (TIM/NKCC^ox^, which is rhythmic under LL). In particular, we outline the necessary steps to analyse two repetitions of the same experiment in which a total of 58 animals were recorded for three to four days in LD and then subjected to constant light for six or seven days. The `metadata.csv` file and all the associated result files can be downloaded on zenodo (https://doi.org/10.5281/zenodo.1172980) [36].

Fist, we install the `rethomics` packages (see the webpage for installation instructions), download the zip archive containing the raw data and extract all files into our working directory. Then, we load the necessary `rethomics` packages:

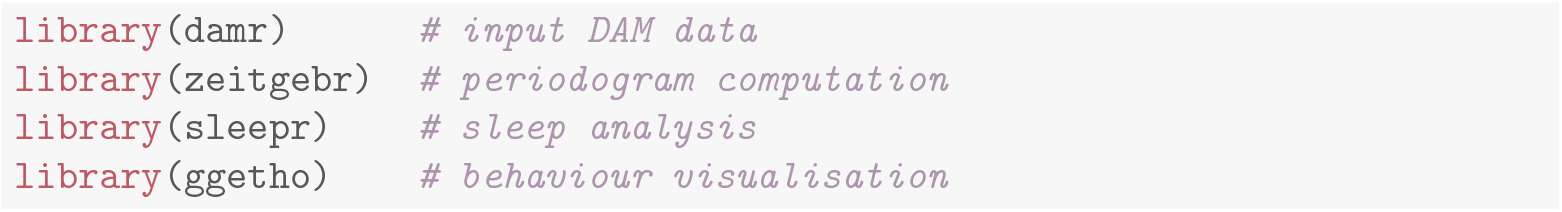

Then, the metadata file is read and linked to the `.txt` result files.

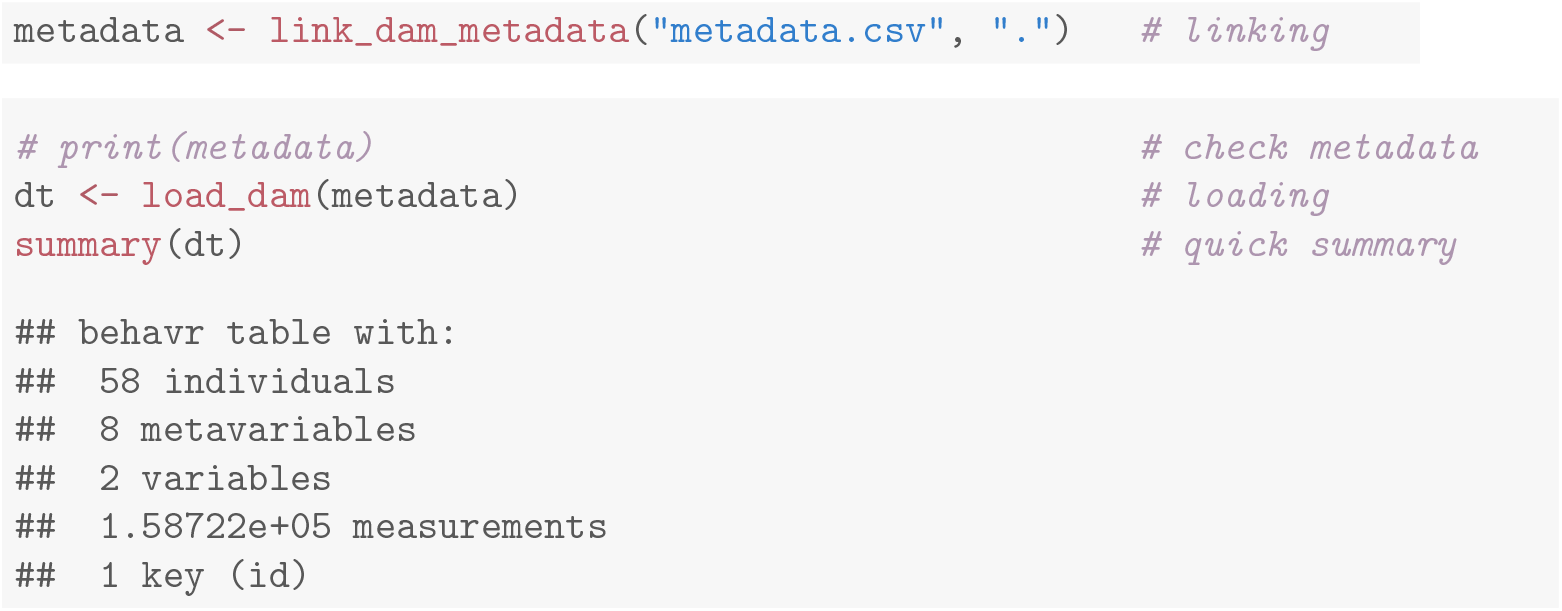

#### Preprocessing

Since the two original replicates do not have the same baseline duration and we want to analyse them together, we align their respective times to the experimental perturbation: the transition from LD to LL (*t* = 0). This is achieved by subtracting the `baseline_days` metavariable from the `t` variable. This gives us an opportunity to illustrate the use `xmv()`, which expands metavariables as variables. In addition, we use the `data.table` syntax to create, in place, a moving variable. It is `TRUE` when `activity` is greater than zero and `FALSE` otherwise:

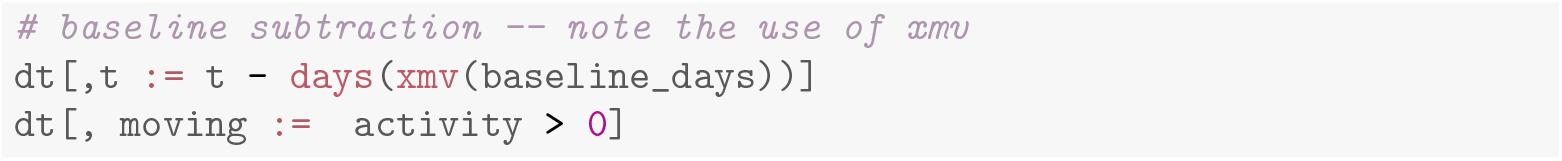

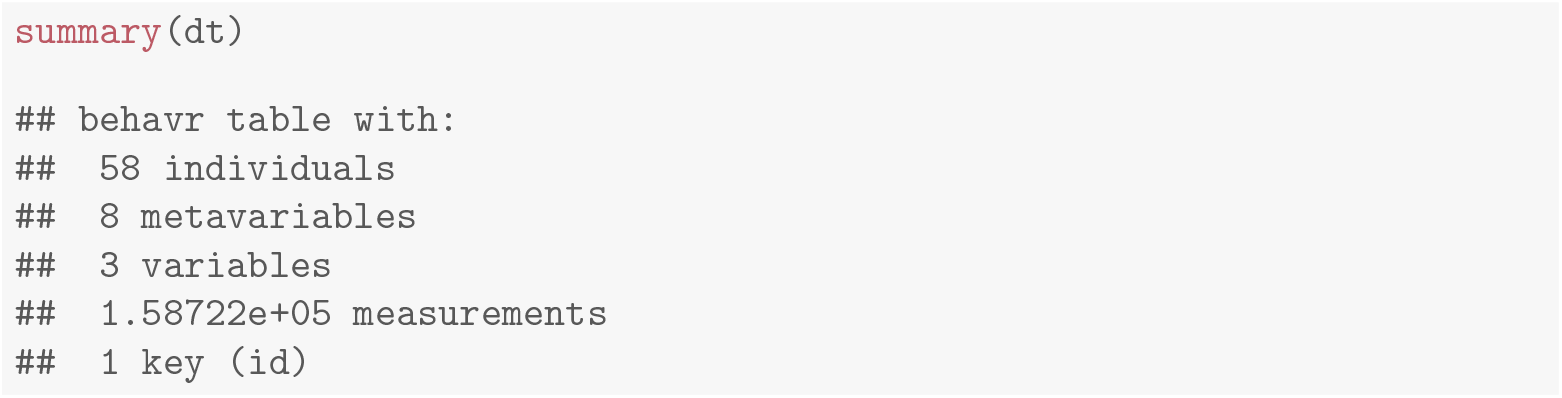

The `id` is a long string of character (for instance,`‘2013-11-19 09:00:00|Monitor36.txt|01’`), which makes it difficult to read and display as a label on a plot. To address this issue, we create our own `label` metavariable, as the combination of a number and `genotype` (e.g. ‘1.NKCCOX/+’). In the restricted context of this analysis, `label` acts as a unique identifier. Importantly, we also retain `id` as an *unambiguous* unique identifier. Indeed, two animals in separate experiments may have the same label, but different `id`s. In addition, if the metadata changes – for instance by the addition or removal of individuals – the label is likely to change, not the `id`.

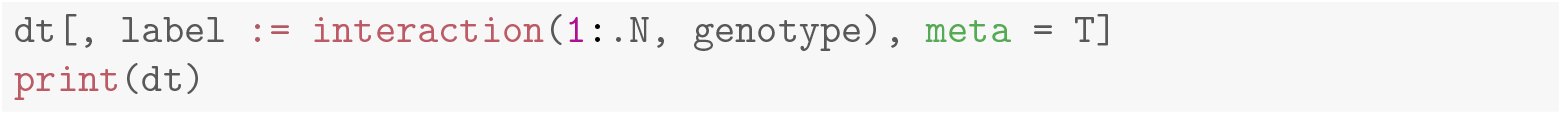

#### Curation

It is important to visualise an overview of how each individual behaved and, if necessary, amend the metadata accordingly. For this, we generate a tile plot (Fig 3A).

**Fig 3.**
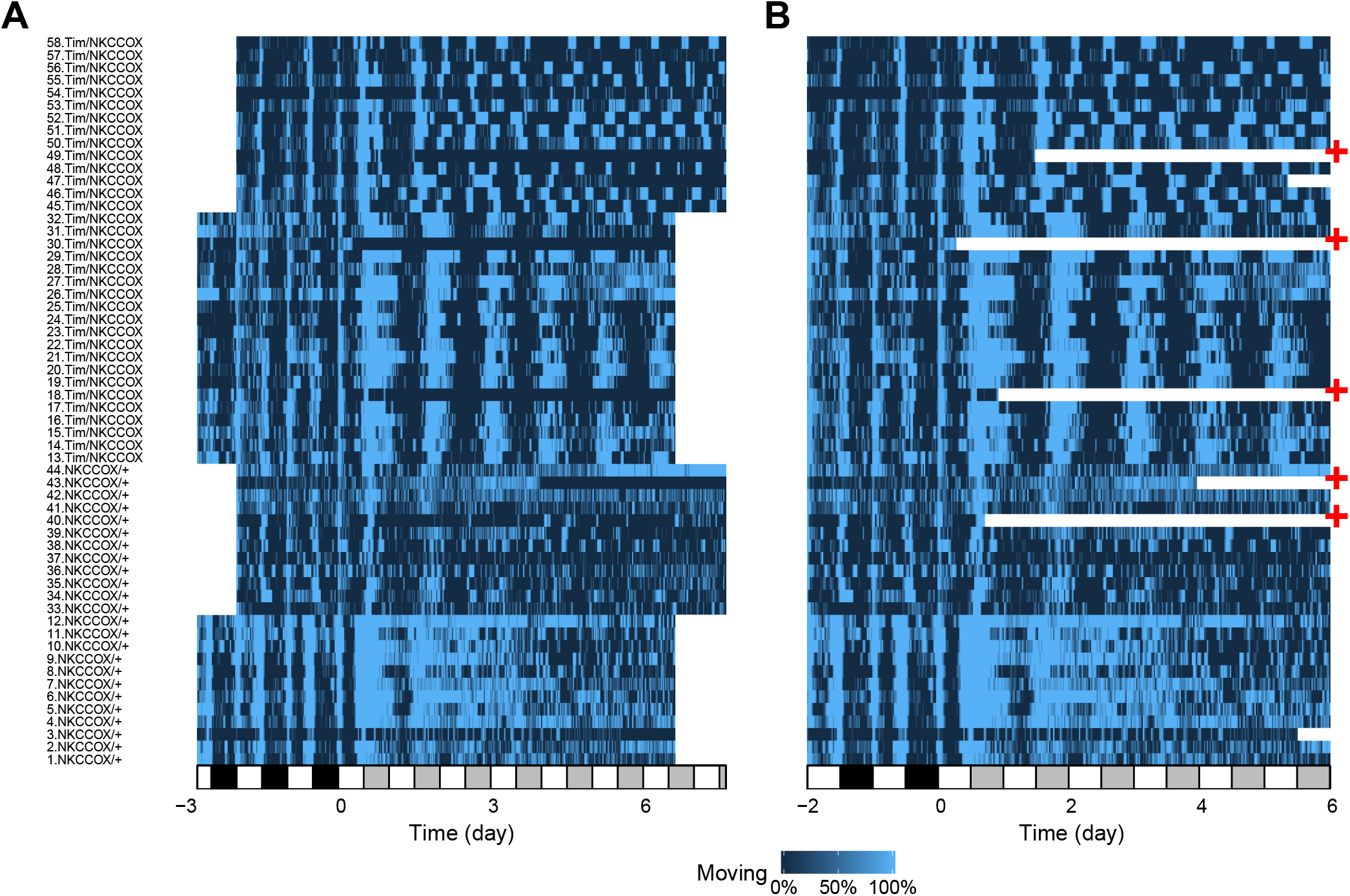
Experiment quality control. Tile plot showing the fraction of time spent moving as a colour intensity. Each individual is represented by a row and time, on the x-axis, is binned in 30 minutes consecutive epochs. A: Uncurated raw data. B: Data after the curation step. Time was trimmed and data from dead animals removed. The red ‘+’ symbols show animals that were removed from the subsequent analysis as they did not survive five complete days in LL.

The white and black rectangles on the time axis show L and D phases, respectively. In the LL regime (for *t* > 0), grey rectangles represent subjective nights.

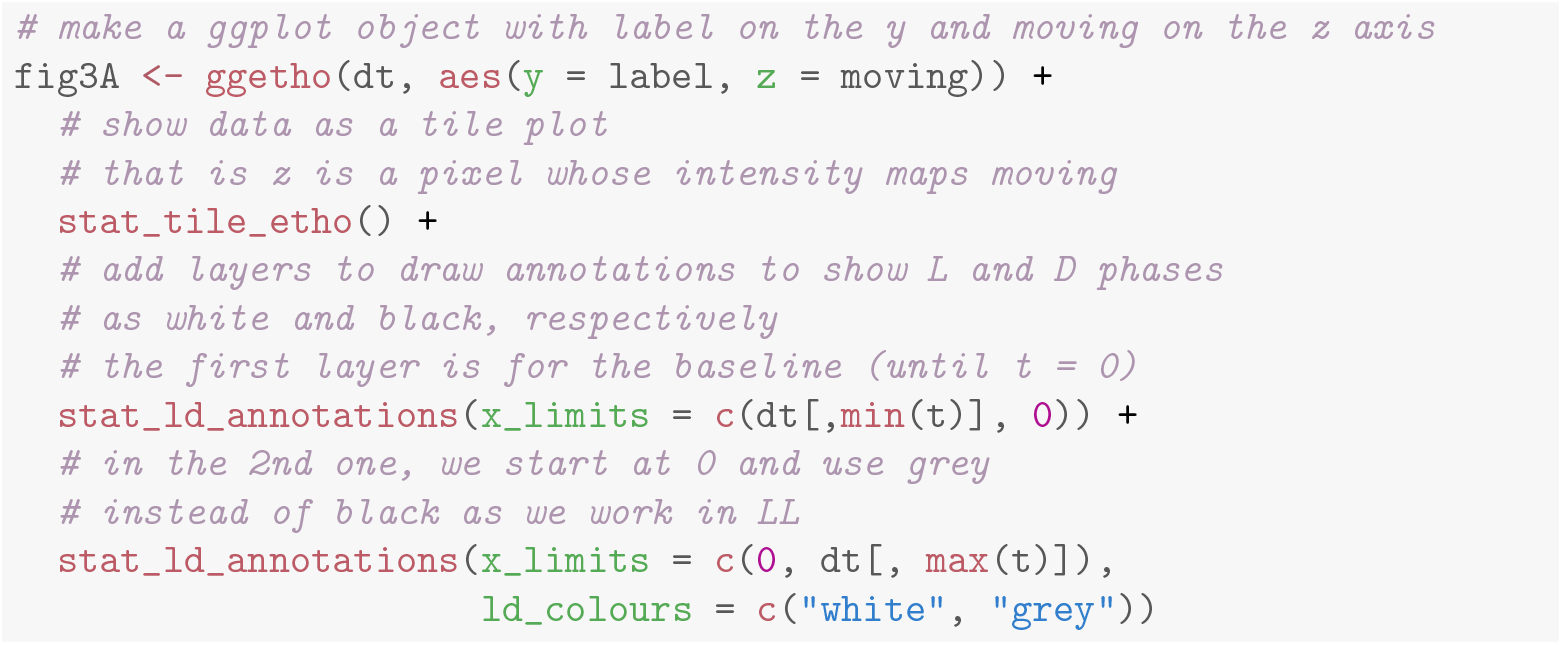

The activity of dead or escaped animals is scored as long series of zeros, which may be erroneously interpreted as inactivity (see, for instance, individual labelled `30` and `18` in Fig 3A). The `sleepr` package offers a tool to detect and remove such data. It proceeds by detecting the first time an animal is immobile for more than 99 % of the time (the default) for at least `time_window` seconds and then discard any subsequent data.

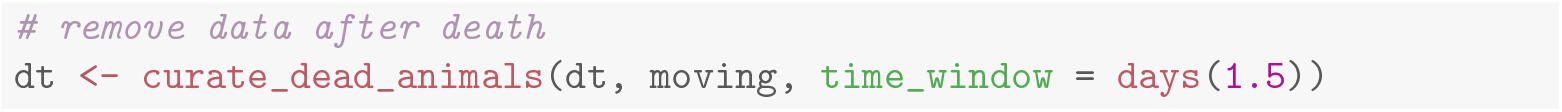

In addition, we can trim the data to the same number of days across experiments and individuals.

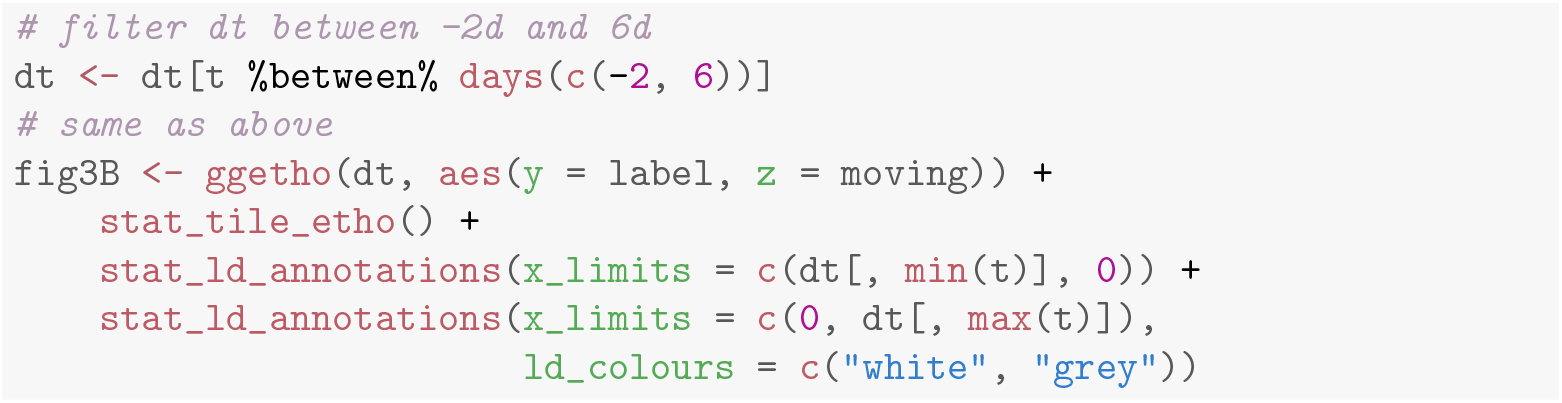

For the purpose of this example, we also exclude animals that died prematurely, and keep only individuals that have lived for at least five days in LL. An overview of the curate data can be visualised in Fig 3B.

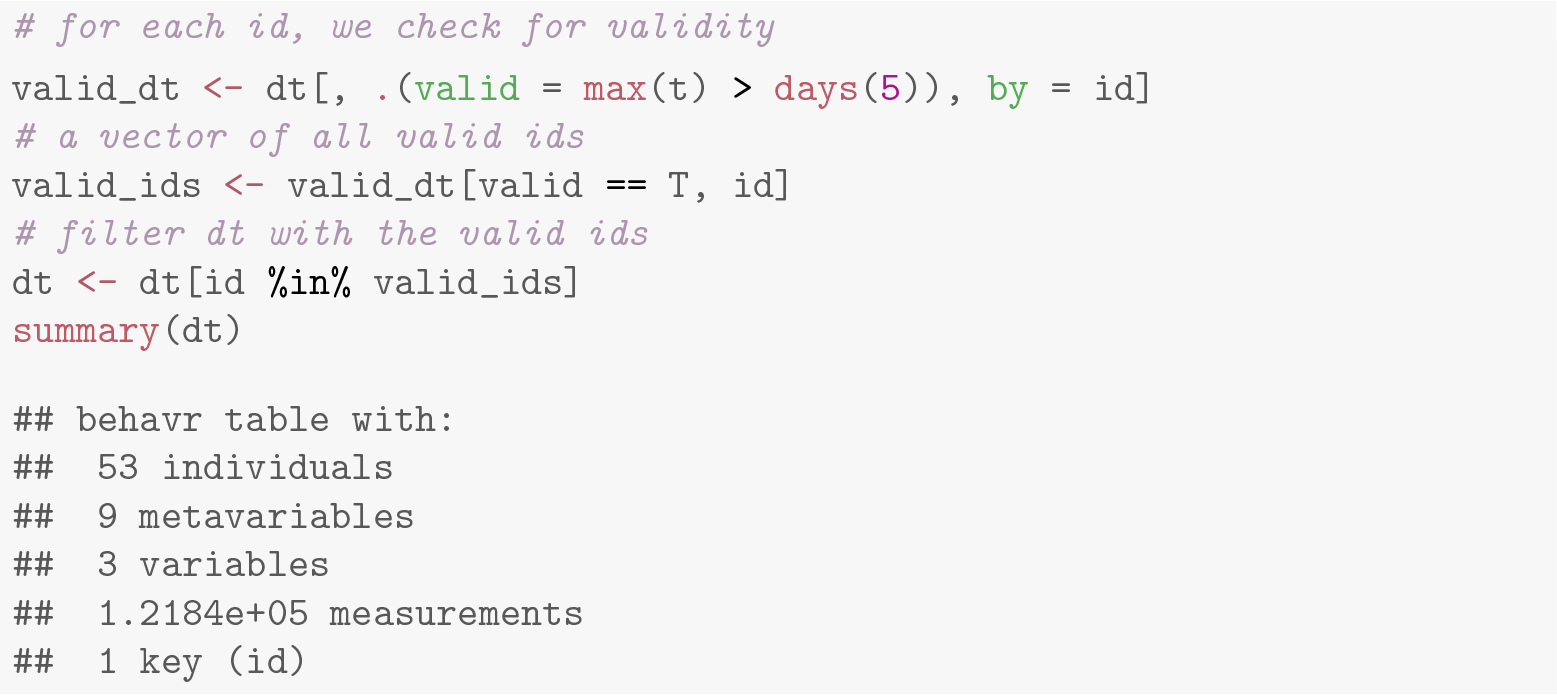

Note that, as a consequence of the curation procedure, we now have 53 ‘valid’ individuals.

#### Double-plotted actograms

Double-plotted actograms are a common choice to visualise the periodicity and rhythmicity in circadian experiments. In S2 FigA, we show a double-plotted actogram for each animal. A selected sample of four individuals for each genotype is shown in Fig 4A.

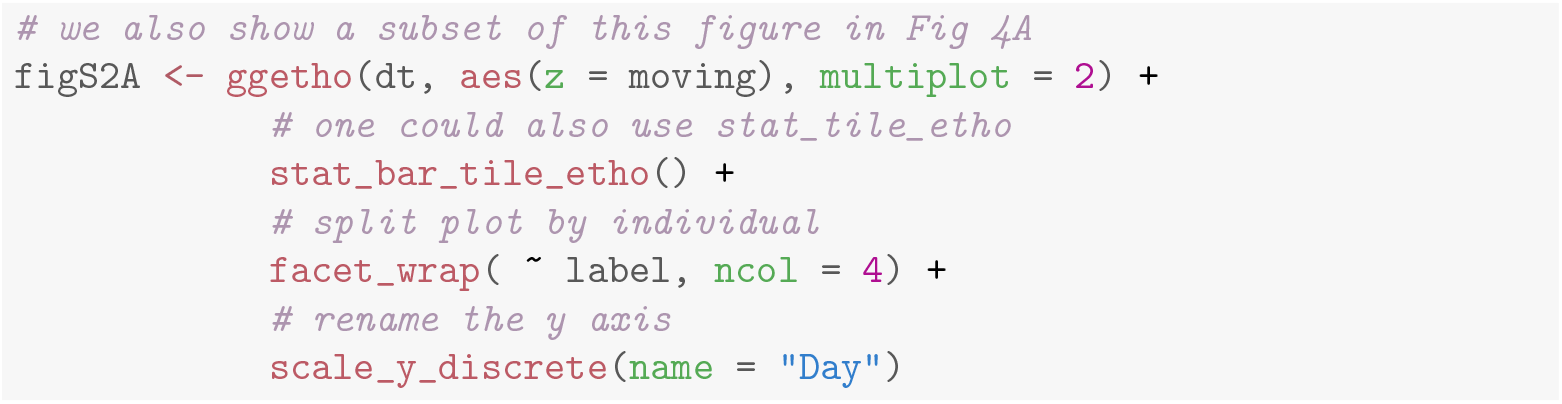

**Fig 4.**
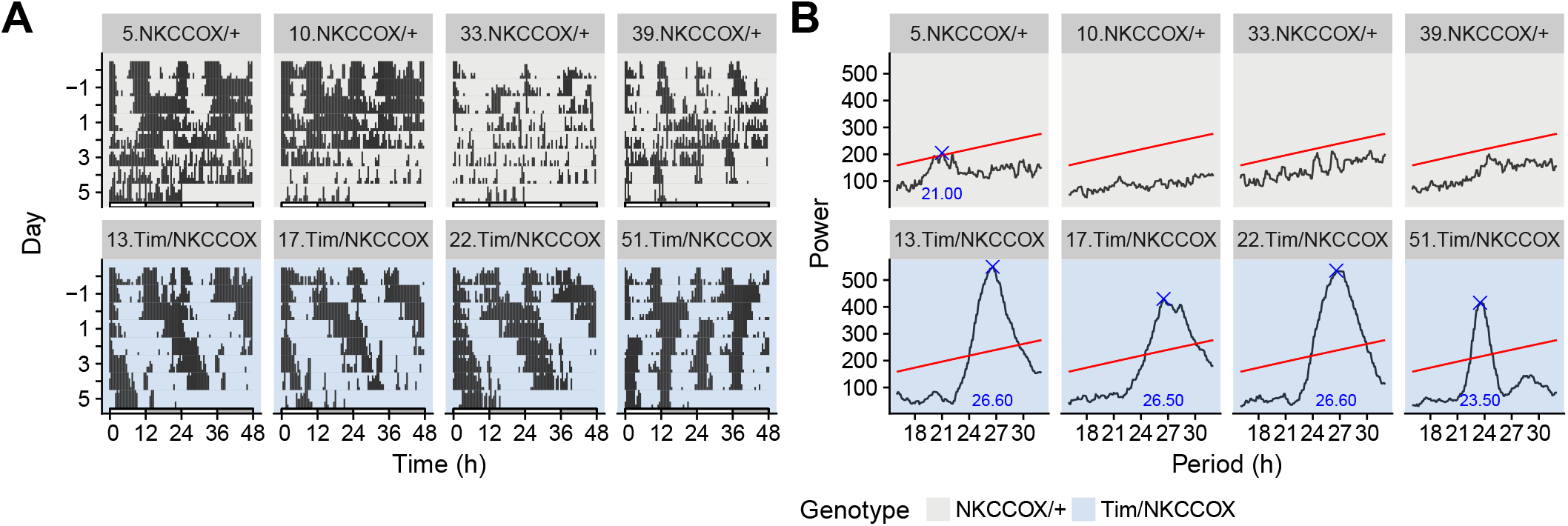
Visualisation of the periodicity in the activity of eight selected animals. A: Double-plotted actograms showing all activity during experiment. Time is defined relative to the transition from LD to LL (at day 0). B: *χ*^2^ periodograms over the LL part of the experiment matching the animals in A. The blue cross represents the highest peak (if present) above the significance threshold, at *α* = 0.05, (red line). Titles on top of each facet refer to the label allocated to each individual. See S2 Fig for all 53 animals.

#### Periodograms

Ultimately, in order to quantify periodicity and rhythmicity, we compute periodograms. Several methods are implemented in `zeitgebr`: *χ*^2^, Lomb-Scargle, autocorrelation and Fourier. In this example, we generate *χ*^2^ periodograms and lay them out in a grid. Periodograms for the subset of eight animals used in Fig 4A are shown in Fig 4B. See S2 FigB for the visualisation of all individuals.

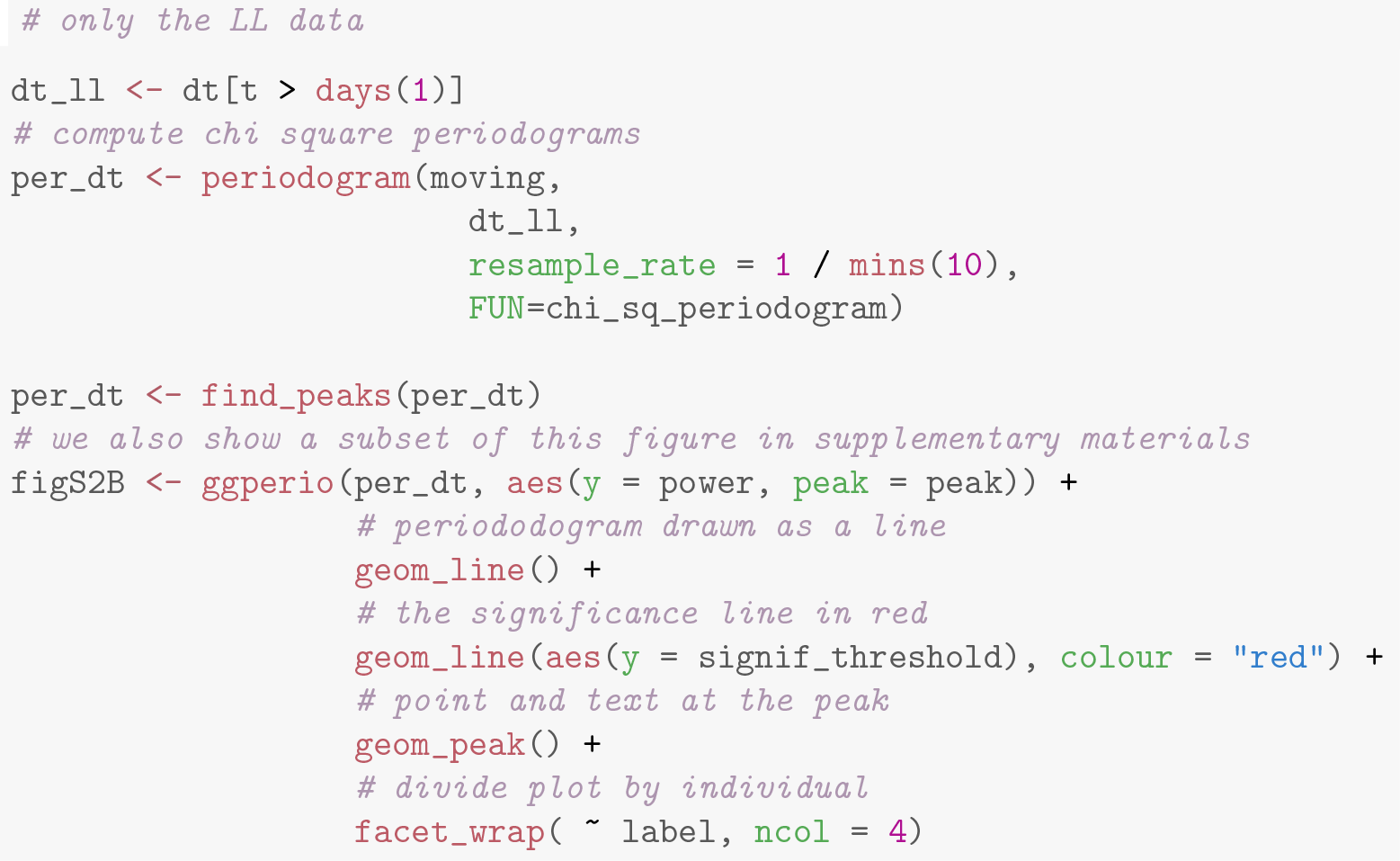

#### Population statistics

As shown in the original study [35], double-plotted actograms and periodograms suggest that NKCC^ox^/+ flies are mostly arhythmic in LL whilst Tim/NKCC^ox^ appear to have a long-period rhythm. To visualise this difference at the population scale, we can plot an average periodogram (see Fig 5A):

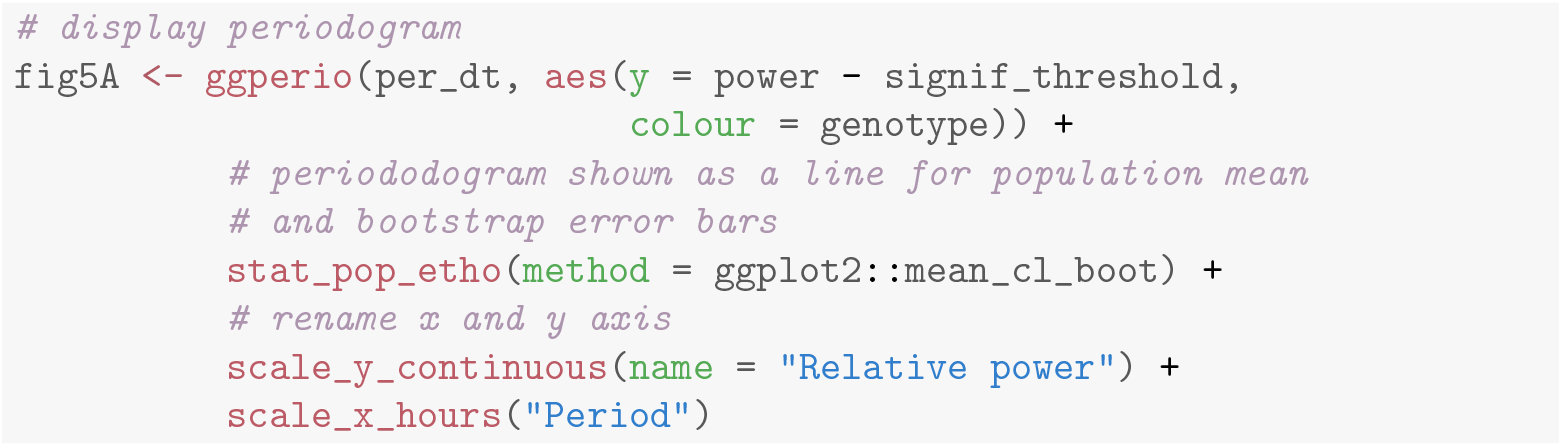

**Fig 5.**
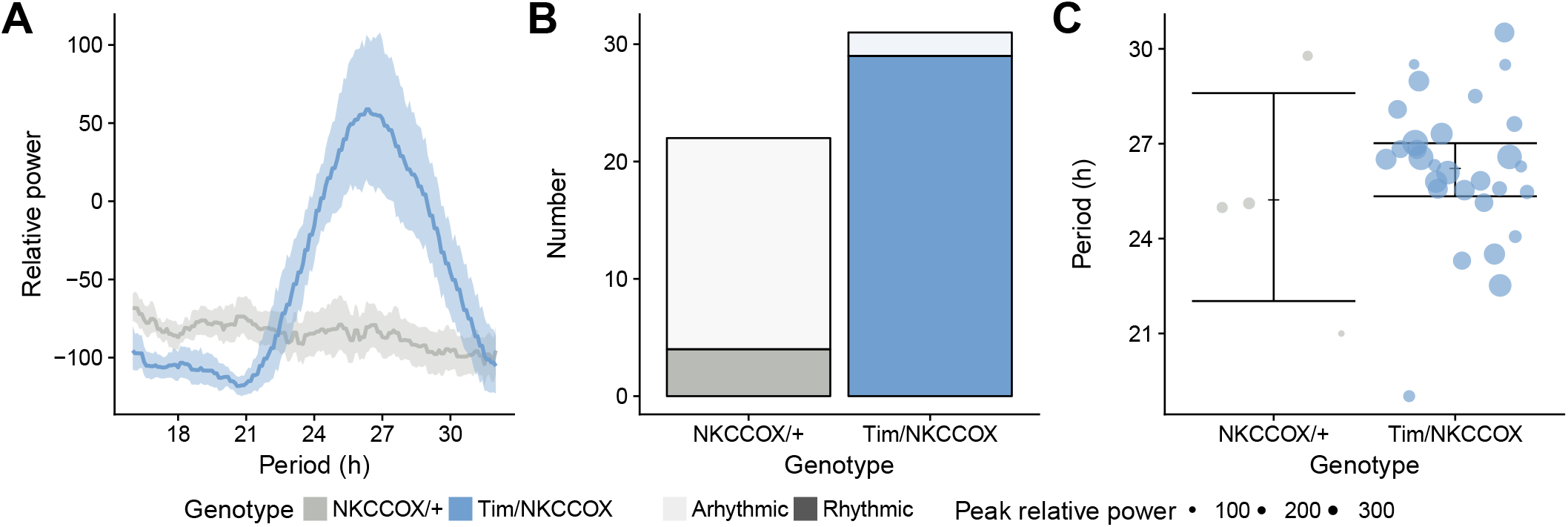
Population statistics on circadian phenotype. A: Average periodograms. The aggregated relative power of the periodogram of all animals. The solid lines and the shaded areas show population means and their 95% bootstrap confidence interval, respectively. B: Frequencies of rhythmic animals. Number of rhythmic animals (*i.e.* with a significant peak) in each genotypes. Dark and clear fillings indicate rhythmic and arhythmic animals, respectively. C: Peak periodicity power and average. Values of the peak period for animals with a peak above the significance threshold, at *α* = 0.05 (*i.e.* rhythmic). Individual animals are shown by dots whose size represent the relative power of the peak period. The error bars are 95% bootstrap confidence interval on the population mean.

To further quantify this difference, we opt to show the number of rhythmic animals – *i.e.* individuals for which a peak was found – in each group (see Fig 5B). Then, we can compare the average value of the peak for the rhythmic animals (see Fig 5C). First of all, we compute a summary per individual (`by=id`):

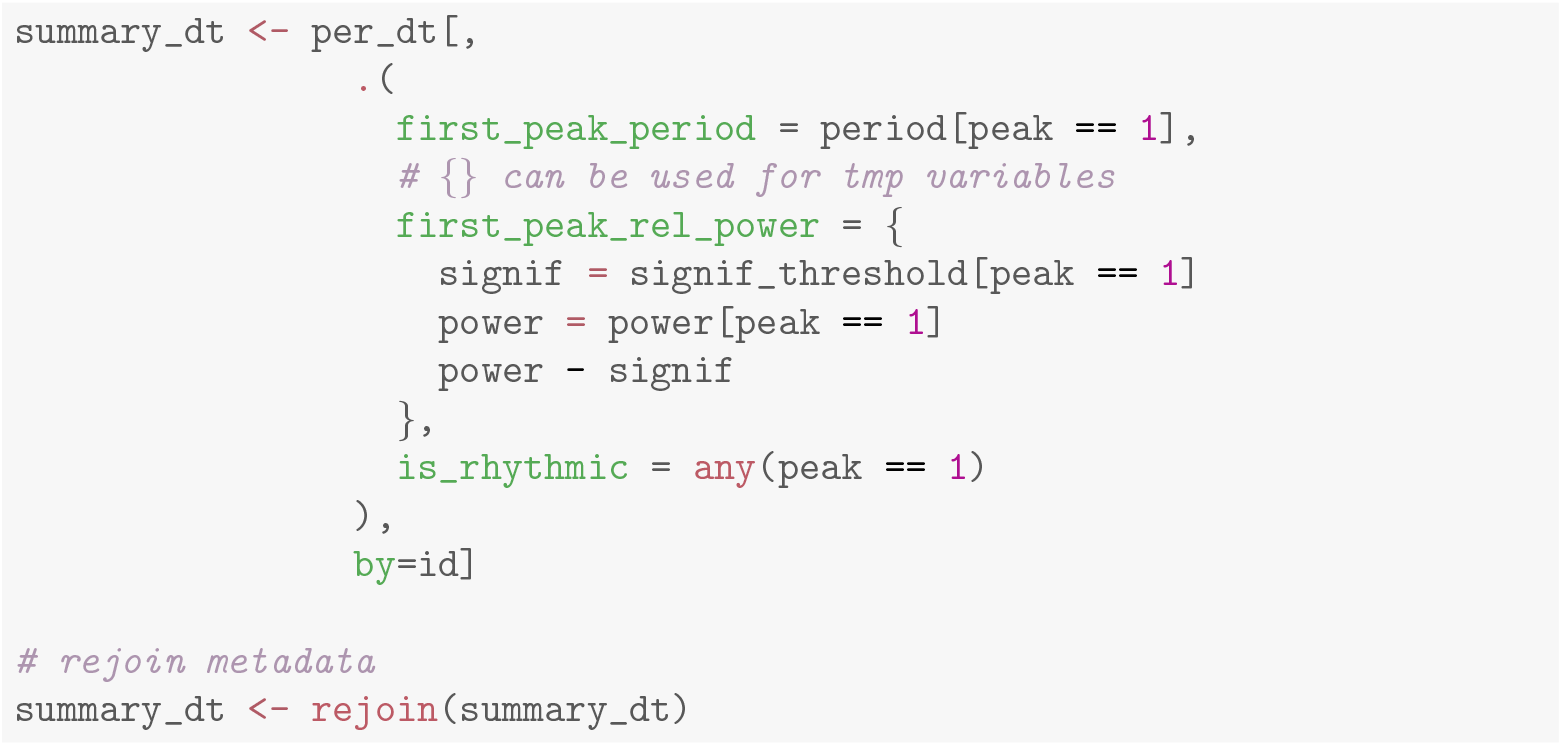

`summary_dt` is a regular data frame with one row per individual, containing both metadata and our summary statistics. It can therefore be used directly by `ggplot`:

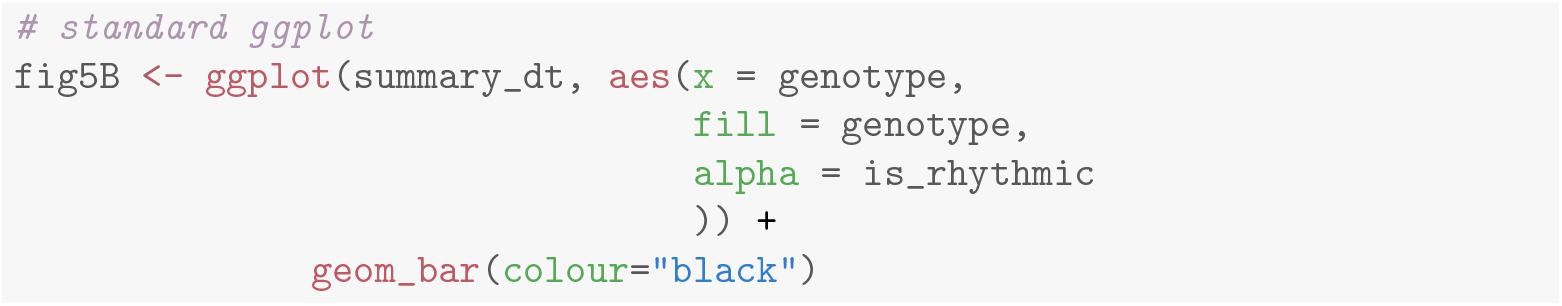

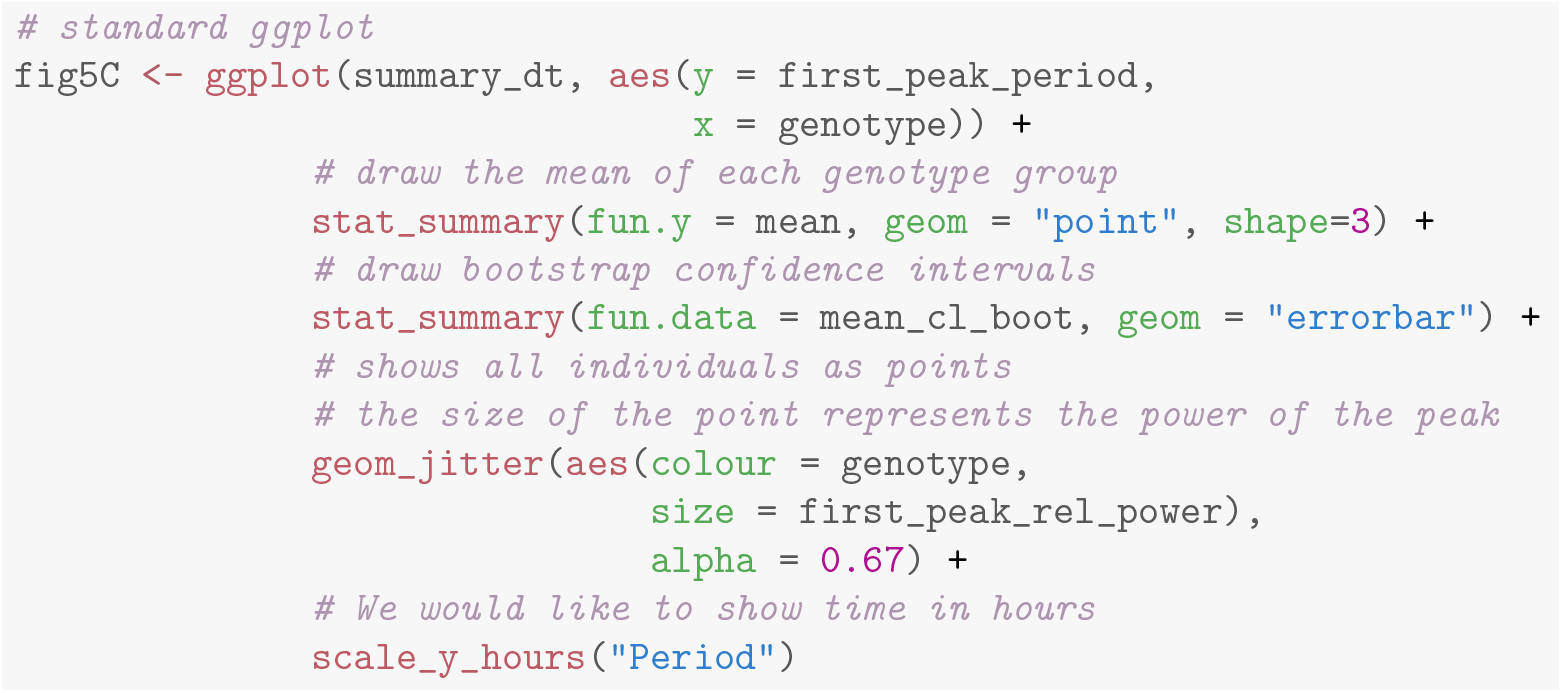

`R` provides one of the richest statistical toolboxes available. Using base `R` we could perform a *χ*^2^ test on the number of rhythmic *vs* arhythmic flies in both genotypes, or, like in this case, fit a binomial generalised linear model:

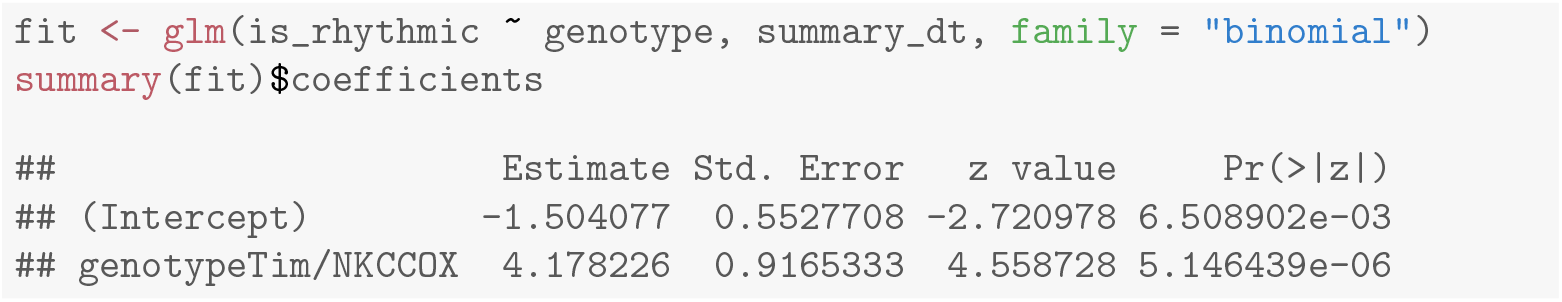

The result shows a strong positive effect of genotype Tim/NKCC^ox^ on the probability of being rhythmic (*p*-value 5.15 × 10^−06^):

Lastly, we can generate a table that compute arbitrary population statistics for each genotype:

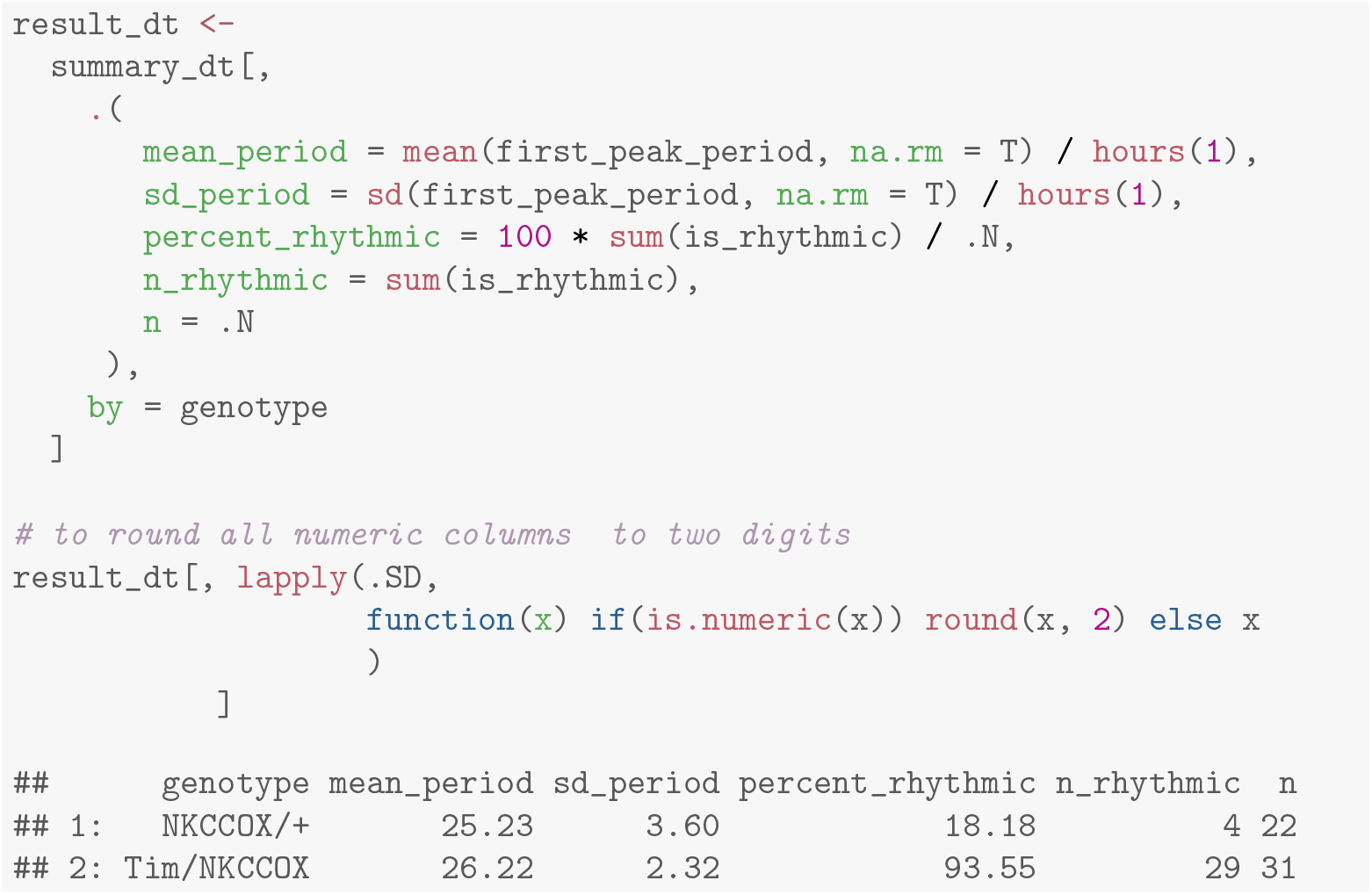

The case study described so far shows how `rethomics` can be employed to generate publication-quality figures and state-of-the-art statistics on pre-existing data. We were able to comprehensively analyse the data from a circadian experiment with a few lines of code, presenting a workflow that applies equally well to much larger datasets.

### Multi-scale analysis of position

One of the challenges of behaviour analysis is the ‘nesting’ of events happening over different timescales. In other words, a behavioural variable can be modulated by the circadian rhythm, but also by co-occurring ultradian and infradian rhythms. For instance, an animal could have rhythmic bursts of locomotor activity recurring at high frequency (*e.g.* 1 min), but also a circadian (*i.e.* 24 h) regulation of the same variable, both rhythms would then happen at timescales separated by approximately three orders of magnitudes, which makes them difficult to visualise and integrate into the same analysis. Being able to keep frequency information over multiple scales is howeverimportant in some cases. In particular, when interested in the frequency modulation of a rhythm by another – that is, if the periodicity of a high-frequency rhythm itself can be a function of a lower frequency one.

The problem of understanding time series at different scales is not uncommon in fields such as economics [37], climate sciences [38] and ecology [39] where variables are governed by multiple underlying rhythms (*e.g.* tidal, daily, yearly and multi-yearly). One approach is to study a variable of interest in the time/period domain using, for instance, continuous wavelet transform (CWT) [40]. In the context of chronobiology, CWT has been suggested as a tool to investigate ultradian rhythms [41].

To illustrate how `rethomic` integrates with other packages and render such non-mainstream analysis possible, we performed a wavelet analysis of the position of 80 naive fruit flies (40 females and 40 males) in their glass tubes. We used the package `scopr`, part of `rethomics`, to load five days of ethoscope positional data, which we sampled at 0.1 Hz. Our variable of interest is the position of animals in their tube (from the food end, *Position* = 0, to the cotton end, *Position* = 1). Fig 6A-C shows the raw position data at two different scales for two representative animals.

**Fig 6.**
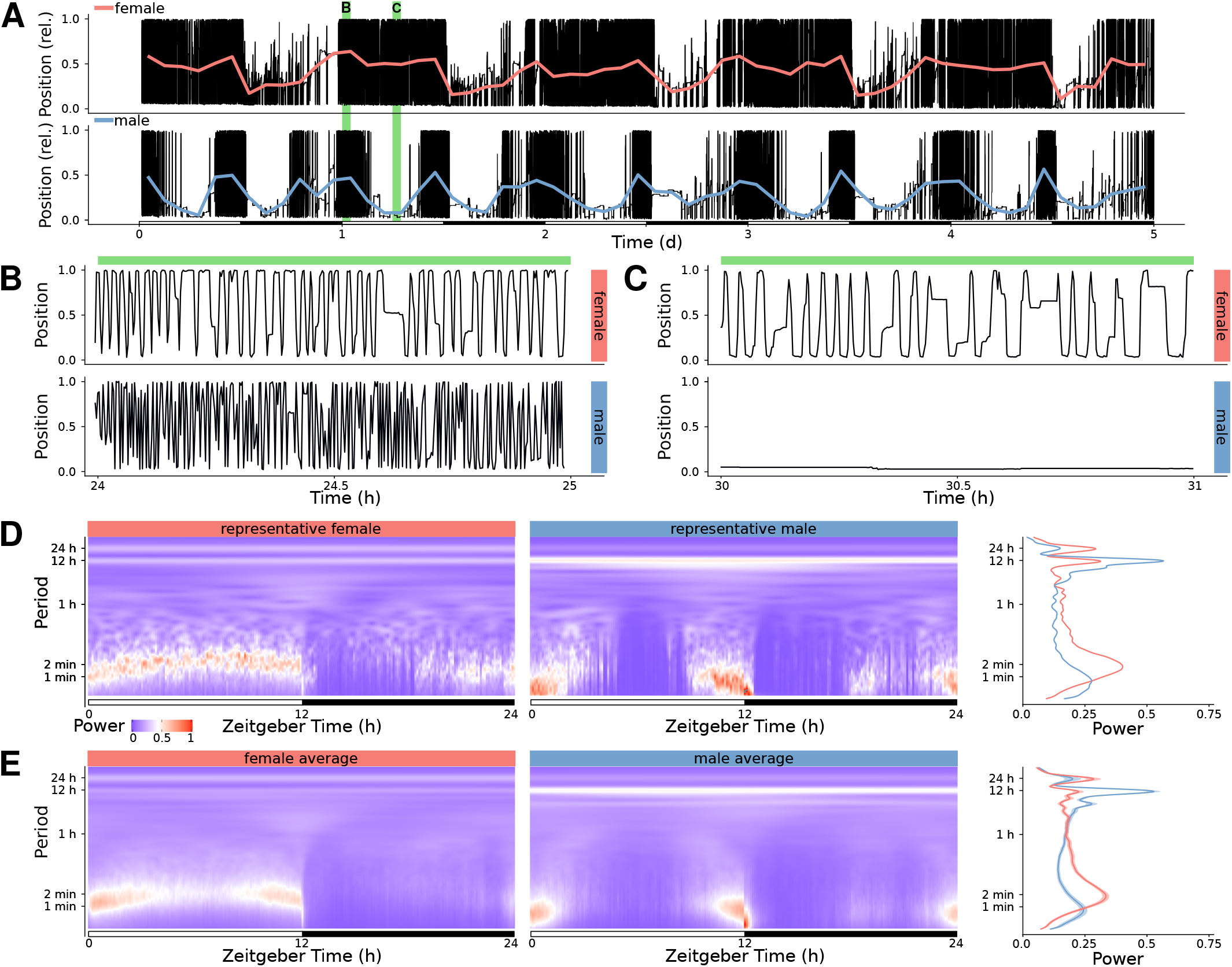
Wavelet analysis of positional data. A: Raw position data for a representative female (top) and male (bottom) *Drosophila melanogaster* over five days, in black. The thick coloured lines show the average position every two hours. The green rectangles in the background show the two time windows selected for B and C. B: Close up of the first time window in A, showing position over one hour, starting at the beginning of the L phase of day 1. C: Close up of the second time window in A, showing position over one hour, starting at the middle of the L phase of day 1. D: Continuous wavelet transform spectrogram for the two representative animals. E: Average spectrogram across 40 males and 40 females. In D and E, the lines on the right show the marginal power spectra corresponding to the shown spectrograms (average across all time). The male data was collected and described in our previous study [21] (controls in figure 5M-P) and the females data was acquired in parallel, in the same experimental conditions, but not previously published.

In order to compute CWT, we used the `WaveletComp` package [42]. We then averaged the result of the five consecutive days in the time/period domain over one circadian day both for the two representative animals (Fig 6D) and for the population(Fig 6E).

As suggested by the slow oscillations of the mean position (Fig 6A), we observe peaks in power corresponding to high-period (12 h and 24 h) rhythms. In addition, a large amount of signal is detected for low-period (around 60 s) events – likely corresponding to the position of animals walking along (back and forward) their tubes in a very paced manner.

Interestingly, in females, this low period pace appears to be frequency modulated during the L phase, suggesting a slower walking speed around ZT6 h. In contrast, males display only a high-frequency rhythm around the phase transitions (L to D and D to L). Surprisingly, the peak of high-frequency rhythm implies a faster pace in males (approximately 60 s) than in females (approximately 120 s) – indicating that, when active, males walk faster than females.

This non-exhaustive proof of principle illustrates how analysis of behavioural data can be taken further by adapting the wide range of numerical tools already available in the `R` ecosystem.

## Availability and Future Directions

All packages in the `rethomics` framework are available under the terms of the GPLv3 license and listed at https://github.com/rethomics/. Extensive installation instructions, as well as reproducible demos and tutorials, are available at https://rethomics.github.io/. All packages are continuously integrated and unit tested on several versions of `R` to minimise the risk of present and future issues.

Several users, in different research groups, have already adopted and are contributing to the future development framework. Several new packages in the `rethomics` framework are currently envisaged. They include utilities to input new behaviour tracking methods and analyse multi-animal interactions.

## Supporting information

**S1 Fig.**
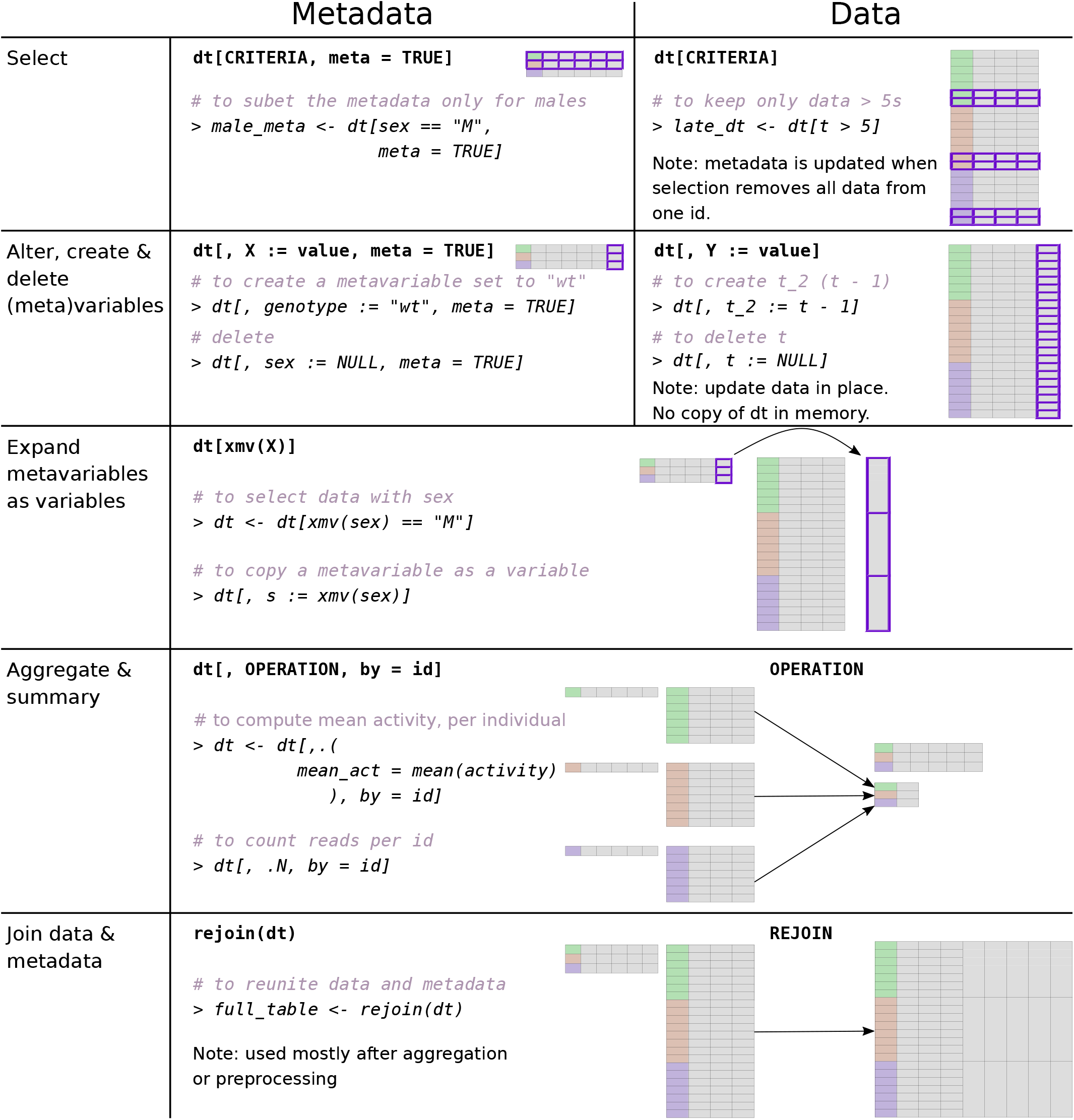
Non-exhaustive list of uses of a behavr table (referred as dt). In addition to operations on data, which are inherited from `data.table`, we provide utilities designed specifically to act on both metadata and data. Commented examples are prefixed by ‘>’.

**S2 Fig.**
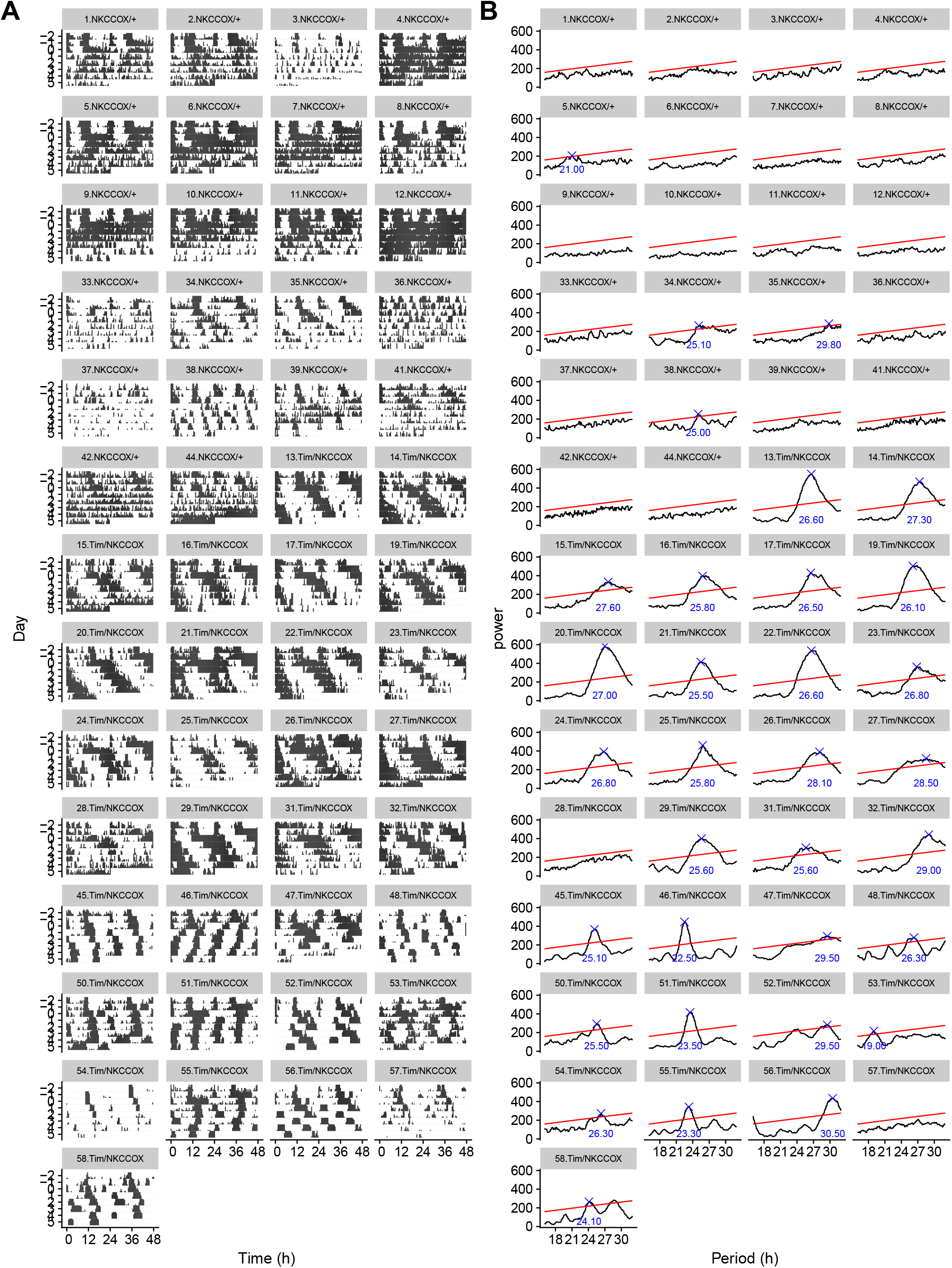
Complete version of Fig 4. See Fig 4 for legend.

## Acknowledgements

We would like to thank people who have directly or indirectly contributed to the this manuscript. In particular, Han Kim, for his invaluable comments on the early versions of `rethomics` and his dedicated contribution to the tutorials; Maite Ogueta and Ralf Stanewsky, for making the DAMS results data available; Alice French, Hannah Jones, Diana Bicazan and Florencia Fernandez-Chiappe for their comments as early users; Marcus Ghosh and Tara Kane for their feedback on the manuscript; Brenna Williams, for her help to support multi-beam DAMS; Patrick Krätschmer, for his time discussing the conceptual framework.

